# Shortcomings of human-in-the-loop optimization for an ankle-foot prosthesis: a case series

**DOI:** 10.1101/2020.10.17.343970

**Authors:** Cara G. Welker, Alexandra S. Voloshina, Vincent L. Chiu, Steven H. Collins

## Abstract

Human-in-the-loop optimization allows for individualized device control based on measured human performance. This technique has been used to produce large reductions in energy expenditure during walking with exoskeletons but has not yet been applied to prosthetic devices. In this series of case studies, we applied human-in-the-loop optimization to the control of an active ankle-foot prosthesis used by participants with unilateral transtibial amputation. We optimized the parameters of five control architectures that captured aspects of successful exoskeletons and commercial prostheses, but none resulted in significantly lower metabolic rate than generic control. In one control architecture, we increased the exposure time per condition by a factor of five, but the optimized controller still resulted in higher metabolic rate. Finally, we optimized for self-reported comfort instead of metabolic rate, but the resulting controller was not preferred. There are several reasons why human-in-the-loop optimization may have failed for people with amputation. Control architecture is an unlikely cause given the variety of controllers tested. The lack of effect likely relates to adaptation protocol or differences in the learning mechanisms or objectives of people with amputation. Future work should investigate these causes to determine whether human-in-the-loop optimization for prostheses could be successful.

## Introduction

Over 600,000 individuals in the United States alone live with a major lower limb amputation, with this number expected to double by 2050 due to the increasing prevalence of diabetes and obesity^1^. Individuals with amputation rely on lower-limb prostheses to replace their lost biological limb. However, gait metrics for users of lower-limb prosthesis are typically worse than those for people without impairment. For example, people using lower-limb prostheses tend to walk more slowly^2^, fall more frequently^3^, and expend more energy to walk at the same speed^4^ when compared to unaffected individuals. In addition, people using a prosthesis tend to demonstrate more gait asymmetry, causing an increased rate of joint degeneration, pain, and osteoarthritis in their intact limb^5, 6^. The combination of all of these factors can lead to limited mobility for individuals with amputation, resulting in secondary health problems, increased medical costs, and more reliance on caregivers^7^.

Although many active ankle-foot prostheses have been developed^8–13^, the vast majority of devices on the market are still passive. Two different approaches in the development of ankle-foot prostheses have had success in reducing the energy cost of walking. The first is to provide ankle power similar to biological gait. Lack of pushoff work of passive prosthesis has been shown to cause higher step-to-step transition losses^14^, which is correlated with increased metabolic rate^15^, although some studies have also shown that more prosthesis push-off work does not necessarily reduce collision work^16,17^. A device that provided pushoff work and emulated biological characteristics of human neuromuscular control, was shown to lead to reductions in the energy cost of walking in people with transtibial amputation by up to 12% compared to passive prostheses^18,19^. The second approach focuses on balance assistance in the form of ankle inversion or eversion torque in response to changes in the frontal plane center of mass. Implementing such control in an ankle-foot prosthesis emulator led to an average 9% reduction in energy expenditure during walking when compared to a zero gain controller^20^. Although these results are promising, simulations demonstrate that active prostheses have more potential to mitigate problems faced by individuals with amputation, with the possibility of reducing the metabolic cost of walking to more than 70% less than unimpaired walking^21^. One possible reason that powered prostheses have not lived up to their potential is because it is still unknown how best to control them, and physiological and neurological differences between users could lead to varying responses to the same device. Previous work has addressed this by hand-tuning control parameters for each subject, but this process is cumbersome and subjective due the high number of parameters that need to be adjusted. Perhaps a control strategy that takes individual gait characteristics into account would allow powered prosthetic devices to better help mitigate problems for individuals with amputation.

Human-in-the-loop optimization (HILO) has been successfully used to determine control parameters for exoskeletons that result in high reductions in metabolic cost^22–26^. This technique involves choosing the parameters of a control architecture based on the human response to changes in control parameters. To do this in a time-efficient manner, HILO uses an optimization strategy that predicts the optimal parameter set over time using a sample of measurements from different control parameter values. Both covariance matrix adaptation evolution strategies (CMA-ES)24and Bayesian optimization^25^ have been successfully used to determine control parameters that lead to reductions in metabolic cost. In one example, optimized assistance from an exoskeleton worn on one ankle reduced the energy cost of walking for all participants significantly more than a hand-tuned static controller, with a range of improvements of 14% to 42% and an average improvement of 24%^24^. HILO has been used to successfully reduce the energy expenditure of running26and inclined walking24with an ankle exoskeleton, in addition to reducing the energy expenditure of walking with a hip exoskeleton^25^. It has also led to reductions in muscle activity while walking with an ankle exoskeleton^24^. These studies suggest that user-specific prosthesis control can provide substantial benefits over conventional, hand-tuned devices. However, HILO has not yet been tested to determine the control parameters of powered prostheses.

In this series of case studies, we applied human-in-the-loop optimization to the control of an active ankle-foot prosthesis used by participants with unilateral transtibial amputation (Fig. 6). Four different classes of control architectures were tested: (1) a heel stiffness controller that varied the stiffness and damping of the heel of the prosthesis (Fig. 1A); (2) a neuromuscular controller that emulated biological components of the muscle-tendon complex and has been previously used to reduce the metabolic cost of walking with an active prosthesis^19^ (Fig. 2A); (3) a balance controller that provided ankle inversion/eversion torque based on deviation of the user’s lateral center of mass velocity and has been previously used to reduce the metabolic cost of walking with a prosthesis emulator20(Fig. 3A); and (4) a time-based torque controller with similar control architecture to that used to reduce the metabolic cost of walking with exoskeletons (Fig. 4A). In addition, both 5-parameter and 4-parameter controllers were implemented for the time-based torque control architecture, resulting in a total of five different control architectures tested.

**Figure 1.**
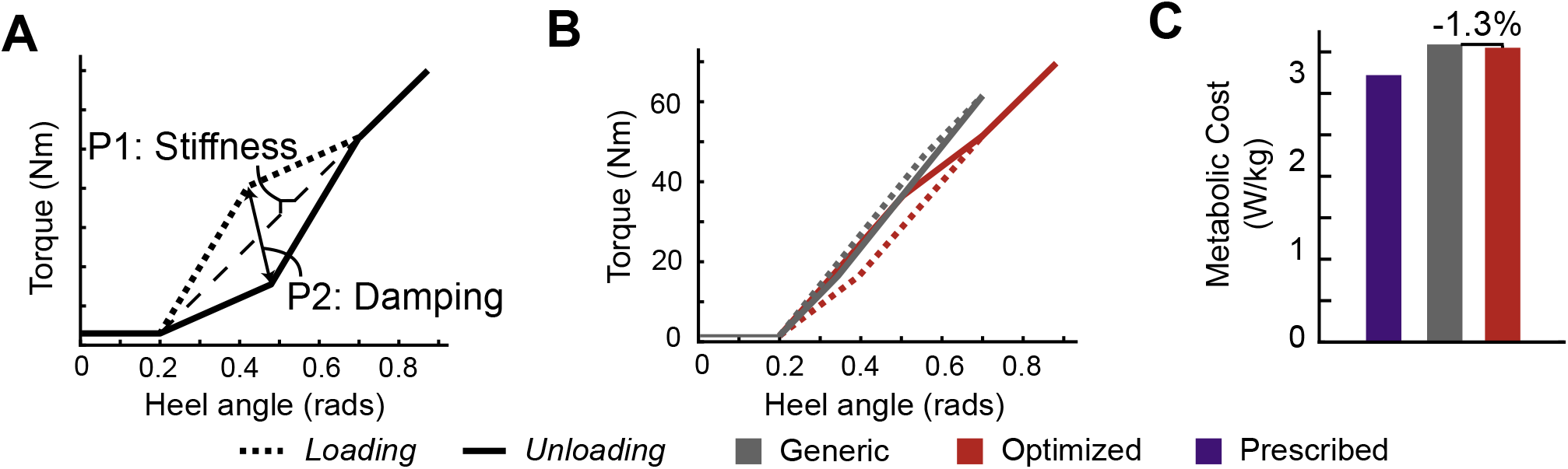
A. The heel stiffness control architecture was dictated by a stiffness and a damping parameter that controlled the torque profile during loading and unloading of the heel as a function of heel angle. Stiffness was allowed to vary between 50 and 130 Nm/rad, and damping from −0.5 to 0.5. B. The resulting torque profile of the generic controller and the optimized controller is shown as a function of heel angle. C. The average metabolic cost of the optimized controller was 1.3% lower than the generic controller, but both were higher than the participant’s prescribed prosthesis.

**Figure 2.**
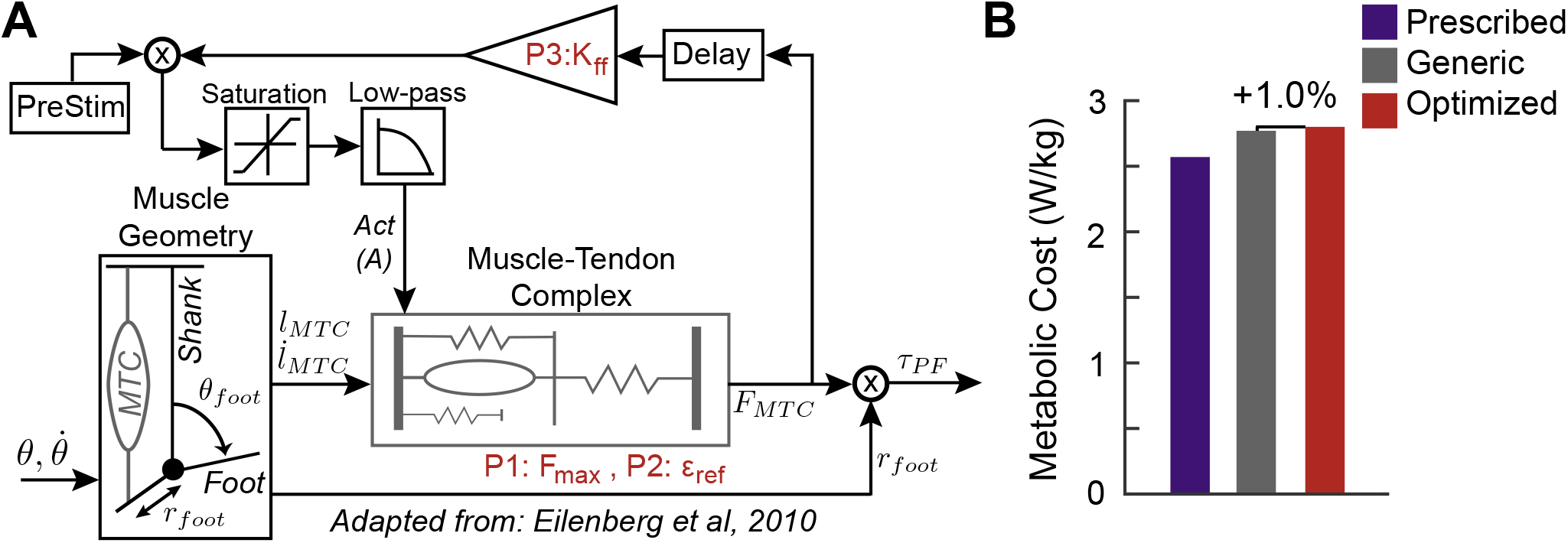
A. The neuromuscular controller draws inspiration from biological muscle-tendon parameters and models the geometry of the prosthetic foot, as well as the the biological muscle-tendon complex. Two optimization parameters *(F_max_* and *ε_ref_*) affect the force output from the muscle-tendon complex model, and the third optimization parameter K_ff_ affects the gain from this force output. Based on biological data, *F_max_* was allowed to vary from 3000 to 10,000 N, and *ε_ref_* from 0.03 to 0.14. Based on prior work, K_ff_ was allowed to vary from 0.7 to 1.5. B. The metabolic results show that the optimized controller was the most costly, followed by the generic controller, and then the prescribed prosthesis.

**Figure 3.**
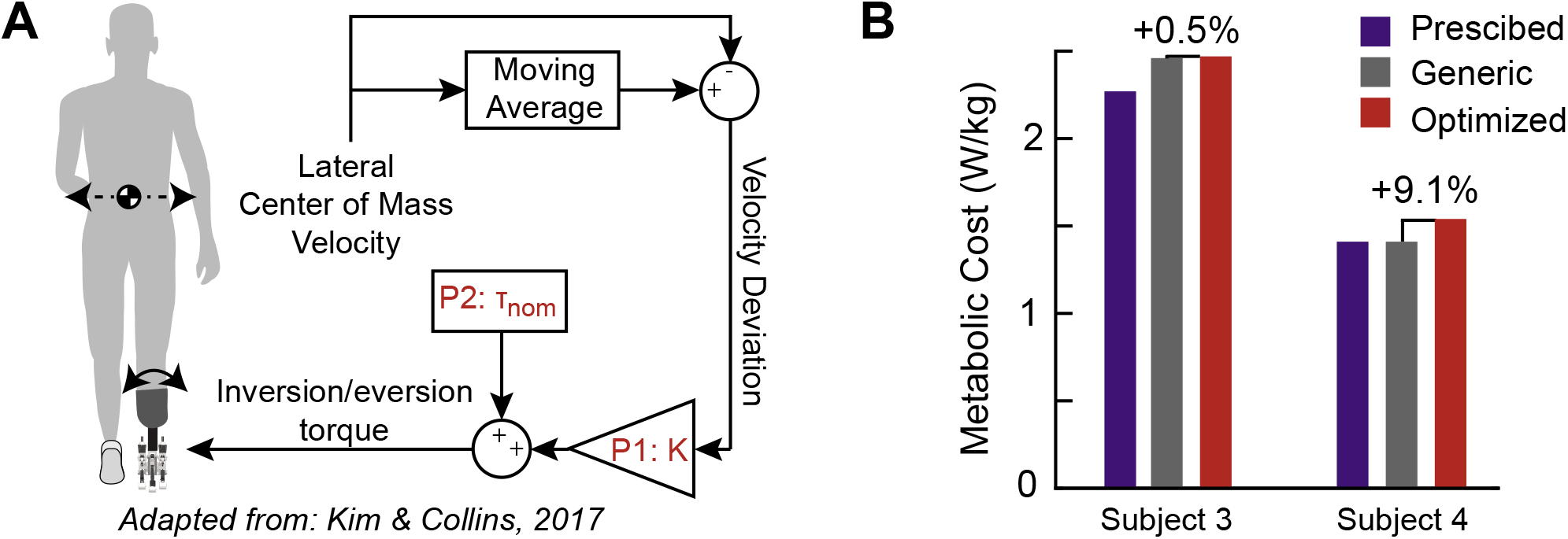
A. The balance controller corrects deviations in center of mass velocity of the user by applying an opposing ankle inversion or eversion torque, scaled by a gain (K), in addition to a baseline nominal torque (τ_nom_). K was allowed to vary from 0 to 5, and *τ_nom_* was allowed to vary from −5 to 5 N·m. B. The optimized controller resulted in higher metabolic cost compared to the generic controller for both participants, with the prescribed prosthesis resulting in a cost equal to or less than the generic controller.

**Figure 4.**
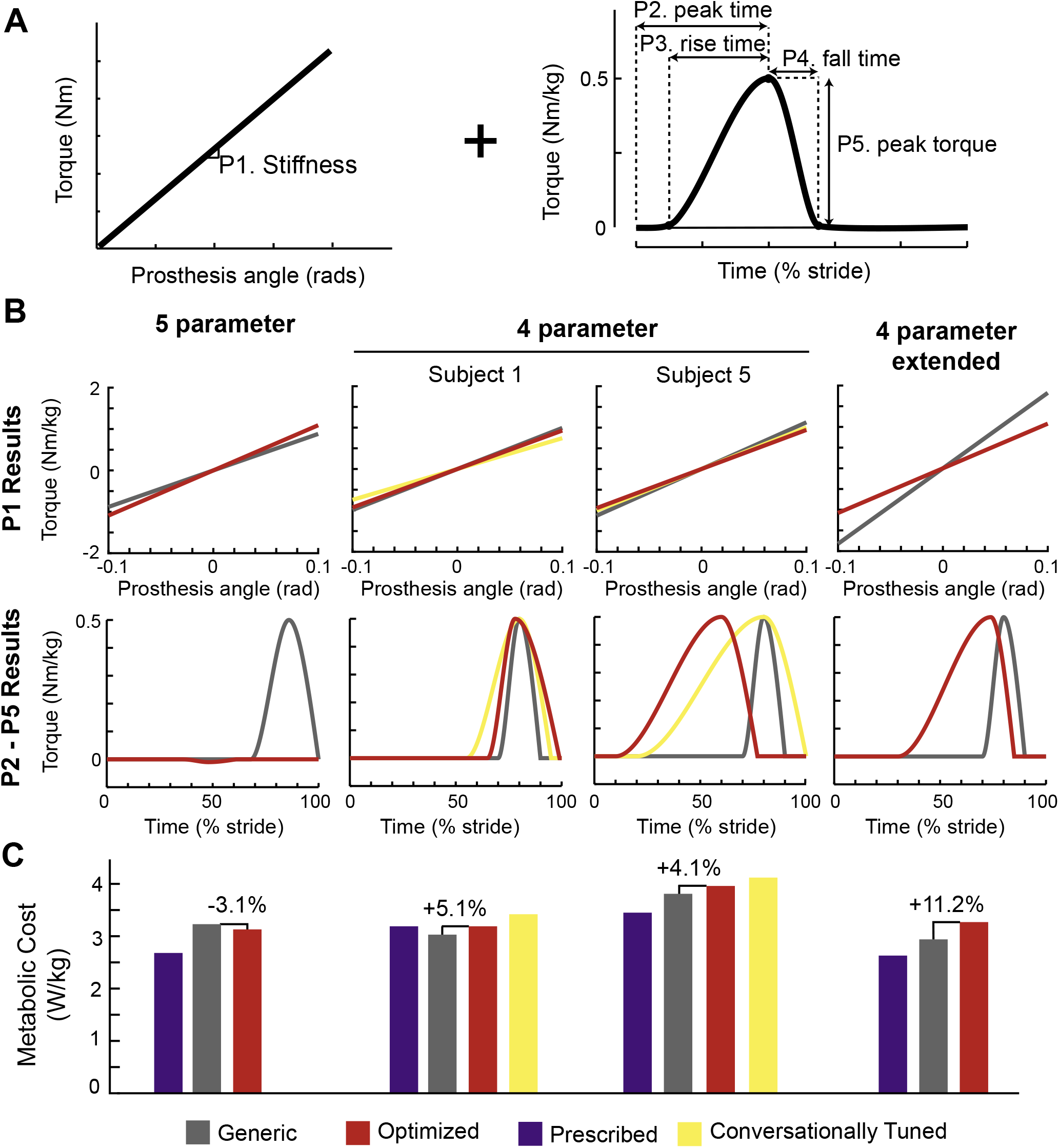
A. The time-based torque controller consisted of an underlying heel stiffness parameter, in addition to three timing parameters (peak time, rise time, and fall time) and one magnitude parameter (peak torque) that affected the shape of the torque profile. Stiffness was allowed to vary from 400 to 2000 N·m/rad, peak time from 10% to 90% of stance, rise time from 10% to 90% of stance, fall time from 10% to 40% of stance, and peak torque from −0.5 to 0.5 N·m/kg. When using the 4-parameter control architecture, peak torque was set at 0.5 N·m/kg. B. The resulting stiffness and torque profiles for each condition in the case studies using the time-based torque controller to minimize metabolic cost is shown. C. Although the optimized controller resulted in a mild reduction in metabolic cost with the 5-parameter control architecture, all other optimized controllers resulted in a metabolic cost higher than the generic controller. The metabolic cost of the conversationally tuned controllers was higher than both the generic and optimized controllers.

**Figure 6.**
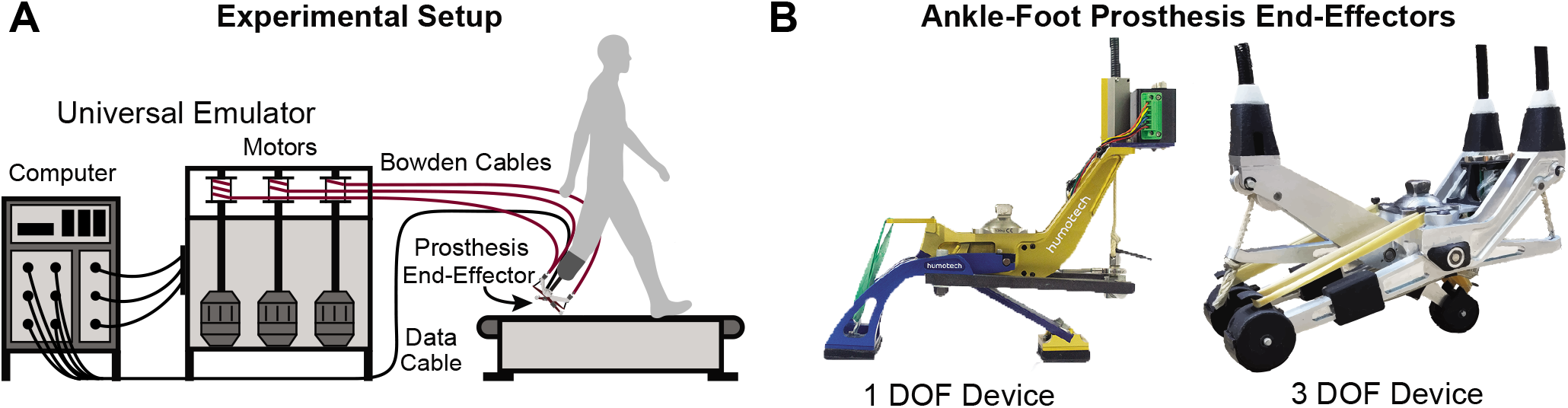
A. The experimental setup used for all case studies is shown. Participants walked on a treadmill in a universal emulator system that consisted of an ankle-foot prosthesis end-effector powered by off-board motors and controlled by a computer. B. Each case study used one of two different ankle-foot prosthesis end-effectors: a one-degree-of-freedom device or a three-degree-of-freedom device.

The main objective of the majority of the case studies was to minimize user energy expenditure. The optimization protocol for each case study with this objective, is outlined in Table 1. Five unique participants were enrolled in these studies, with some individuals completing more than one experiment (Table 2). The protocol was similar for all experiments and was based on an optimization protocol successfully used with exoskeletons^24^. However, there was one exception where the participant completed an extended protocol to determine its effects on adaptation and the optimization. Finally, one additional case study optimized the control parameters with the objective of maximizing participant preference instead of minimizing metabolic rate. For all case studies, participants completed separate validation trials after optimization was complete, and performance with the optimized controller was compared to a generic controller and the participant’s prescribed prosthesis.

**Table 1.**
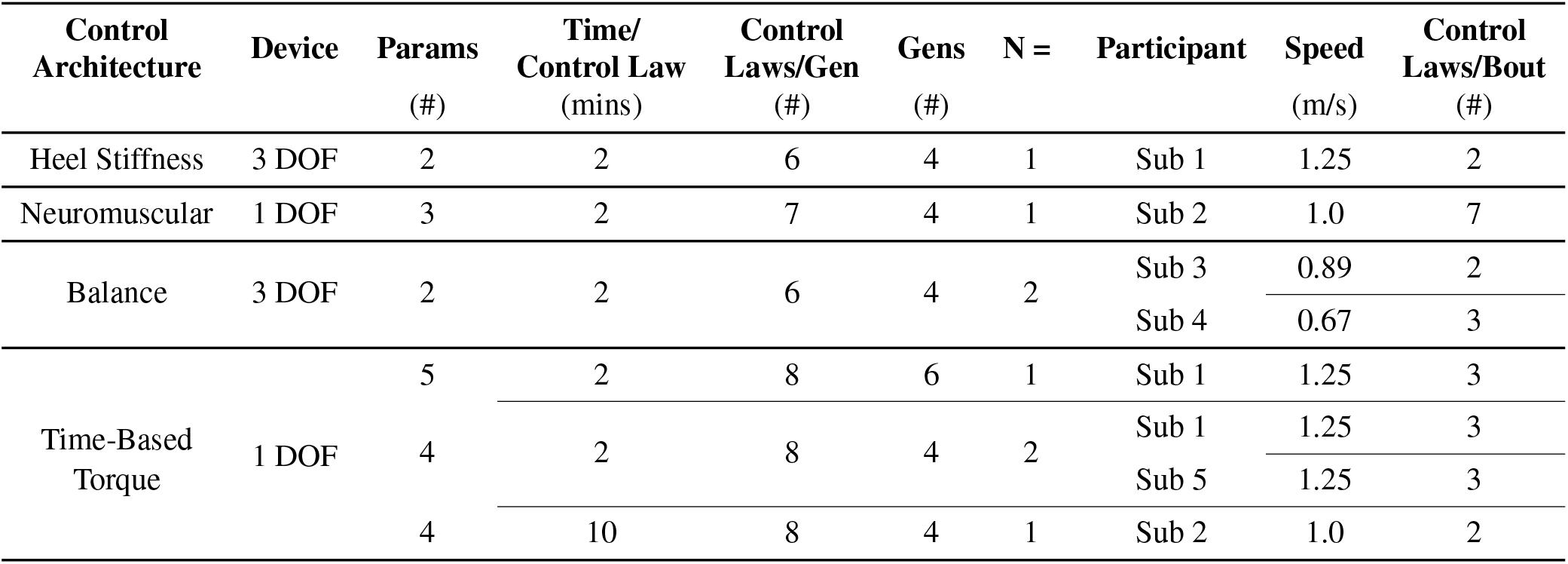
An overview of the protocol for each human-in-the-loop optimization case study that attempted to minimize metabolic rate, including the control architecture and device used, the number of parameters (params) in the control architecture, the time spent walking in each control law with a specific parameter set, and the number of control laws per generation (gen). The participants in each case study are also listed, along with their self-selected walking speed and the number of different control laws that were tested during each bout of continuous walking.

**Table 2.**
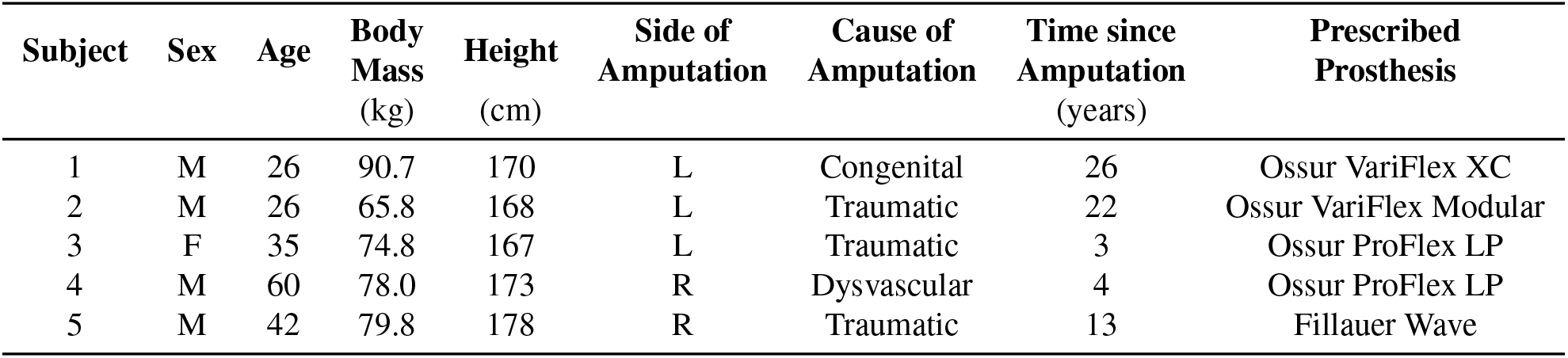
Demographics for the five subjects with unilateral transtibial amputation that participated in the case studies. Note that some participants took part in more than one case study, as described in Table 1.

## Results

### Heel Stiffness Controller

The resulting optimized parameters after four generations of walking were a heel stiffness of 100 N·m/rad with a heel damping coefficient of 0.4. In validation, the optimized controller was compared to a generic controller (stiffness = 120 N·m, damping = −0.5) chosen to provide 32% heel energy dissipation, emulating common energy storage and return (ESR) passive devices on the market^27^. The optimized controller resulted in a 1.3% reduction in metabolic cost compared to the generic controller (optimized: 3.55 W/kg, generic: 3.59 W/kg) (Fig. 1B). Both resulted in higher metabolic cost than walking with the participant’s prescribed prosthesis (3.22 W/kg), which was only tested once in validation for this case study.

### Neuromuscular Controller

Three parameters in the neuromuscular controller were optimized: two within the muscle-tendon complex model *(F_max_* and **ε_ref_**) and one feedforward gain *Kff* (See Fig. 2). After four generations of walking, the optimized parameters for the neuromuscular controller were as follows: *F_max_*= 4261 N, Kff = 1.346, and *ε_ref_* = 0.062. This optimized controller was compared to a generic controller using parameters previously determined to best mimic biological ankle torque *(F_max_*= 3377 N, Kff = 1.22, and *ε_ref_* = 0.04)^19^. In validation, the optimized controller resulted in a metabolic cost that was 1.0% higher than the generic controller (optimized: 2.80 W/kg, generic: 2.77 W/kg) (Fig. 2B). However, both controllers resulted in higher metabolic cost than walking with the participant’s prescribed prosthesis (2.57 W/kg).

### Balance Controller

Two participants (Subject 3 and Subject 4) completed the optimization protocol for the 2-parameter the balance controller consisting of four generations of walking (See Fig. 3A). The optimization for Subject 3 resulted in a K value of 1.24 and a *τ_nom_* value of −0.173, while the optimization for Subject 4 resulted in a K value of 3.97 and a *τ_nom_* value of 3.03. These were compared to a zero gain generic controller where the K value was set to zero, but the optimized *τ_nom_* value was retained. The optimized controller resulted in a 0.5% increase in metabolic cost for Subject 3 (optimized = 2.47 W/kg, generic = 2.46 W/kg) and a 9.1% increase in metabolic cost for Subject 4 (optimized = 1.54 W/kg, generic = 1.41 W/kg). The prescribed prosthesis cost for Subject 3 was less than either controller (2.27 W/kg), while the prescribed prosthesis cost for Subject 4 was equivalent to the generic controller (1.41 W/kg). The results for all validation conditions are shown in Fig. 3B.

### Time-based Torque Controller

#### 5-parameter

The five parameters optimized in this control architecture were: stiffness, peak time, rise time, fall time, and peak torque (Fig. 4A). Because of the increased number of parameters, the participant completed six generations of optimization before validation. His optimal parameters were as follows: stiffness = 990 N·m/rad, peak time = 48%, rise time = 12%, fall time = 13%, peak torque = −0.01 N·m/kg. Because the optimal peak torque value was so small in magnitude, it should be noted that varying the peak time, rise time, and fall time had minimal effects on the resulting torque profile. This optimized controller was compared in validation with a generic controller with the following parameters: stiffness = 800 N·m/rad, peak time = 86%, rise time = 18%, fall time = 14%, and peak torque = 0.05 N·m/kg. The optimized controller resulted in a 3.1% decrease in metabolic cost from the generic controller (optimized = 3.13 W/kg, generic = 3.23 W/kg). However, both controllers resulted in higher metabolic cost than the participant’s prescribed prosthesis (2.68 W/kg). Visualization of the torque profiles of the controllers and resulting metabolic cost from all validation conditions are shown in Fig. 4B and Fig. 4C.

#### 4-parameter

Because the peak torque chosen by the optimizer for the 5-parameter time-based torque control optimization had a magnitude so close to zero, negating the effect of the other parameters, additional experiments were run on a 4-parameter time-based torque control architecture where the peak torque was set at 0.5 N·m/kg. Two participants (Subject 1 and Subject 5) completed the optimization protocol for this control architecture. Subject 1 had previous experience walking in the 5-parameter control architecture, and his resulting optimal control parameters after four generations were as follows: stiffness = 843 N·m/rad, peak time = 78%, rise time = 13%, fall time = 21%. This was the first case study that Subject 5 had participated in, and his resulting optimal control parameters were as follows: stiffness = 754 N·m/rad, peak time = 60%, rise time = 50%, fall time = 17%. These optimized controllers were compared to a generic baseline controller with the following parameters: stiffness = 900 N·m/rad, peak time = 80%, rise time = 10%, fall time = 10%. In validation, the optimized controller resulted in a 5.1% increase in metabolic cost for Subject 1 (optimized = 3.19 W/kg, generic = 3.03 W/kg) and a 4.1% increase in metabolic cost for Subject 5 (optimized = 3.96 W/kg, generic = 3.81 W/kg). The prescribed prosthesis was also tested in validation. Subject 1’s prescribed prosthesis resulted in the same metabolic rate as the optimized controller (3.19 W/kg), while Subject 5 had a more typical lower metabolic rate with his prescribed prosthesis (3.45 W/kg). The torque profiles and average metabolic results of the validation trials are shown in Fig. 4B and Fig. 4C.

#### Conversationally Tuned

During the validation of the 4-parameter optimization using the time-based torque control architecture, a fourth condition based on subject preference was tested in addition to the prescribed prosthesis, generic controller, and optimized controller. To determine the parameters of the controller for this fourth condition, the subject walked on the prosthesis as the controller was changed and gave verbal feedback on the preferred parameters. The preferred parameters chosen by Subject 1 were: stiffness = 675 Nm/rad, peak time = 80%, rise time =25%, fall time = 15%. The preferred parameters chosen by Subject 5 were: stiffness = 800 Nm/rad, peak time = 80%, rise time = 60%, fall time = 20% (igure4B). In validation, the conversationally tuned controllers resulted in higher metabolic rate than all other conditions for both Subject 1 (3.42 W/kg) and Subject 5 (4.12 W/kg) (Fig. 4C). We also collected each subject’s opinions of the controllers during validation. Subject 1 commented that the generic controller was “comfortable from the start and stayed comfortable,” but he also noted that it “has finicky timing.” He noted that both the optimized controller and the conversationally tuned controller were “comfortable from the start,” but the optimized controller “felt like it required more effort later” and the conversationally tuned controller “didn’t feel good later.” Subject 5 noted that the generic controller had “too much pushoff,” the conversationally-tuned controller had “definitely not enough pushoff,” and the optimized controller had “a perfect amount of pushoff.”

#### Extended protocol

One subject completed an optimization protocol with the 4-parameter time-based torque controller in which the time spent in each control law was increased by a factor of five, from two minutes to ten minutes, in order to give the participant more time to adapt to each controller. The optimization resulted in the following control parameters: stiffness = 710 Nm/rad, peak time = 74%, rise time = 44%, fall time = 11%. The optimal controller was compared to a generic controller with the following parameters: stiffness = 1200 Nm/rad, peak time = 80%, rise time = 10%, fall time = 10% (Fig. 4B). The optimized controller resulted in an 11.2% increase in metabolic cost compared to the generic controller (optimized = 3.27 W/kg, generic = 2.94 W/kg). Both resulted in higher cost than the participant’s prescribed prosthesis (2.63 W/kg; Fig. 4C).

To examine if the additional time spent in each control law resulted large changes in metabolic rate between the conditions, we compared the results of a 2-minute metabolic fit, as performed in all other case studies, with the metabolic results averaged in the last two minutes of each ten-minute control law. The RMSE error between these two conditions was 0.24 ±0.16 W/kg, or 6% of the participant’s metabolic cost while walking with the prescribed prosthesis (without subtracting the standing baseline, as this was only collected during optimization and not validation). Because the optimizer uses a weighted average of the control laws resulting in the lowest metabolic cost from each generation to create the distribution of the parameter set for the next generation, we also compared the rankings using the 2-minute metabolic fits to the rankings using the average of the last two minutes of each condition. In all generations, the 2-minute metabolic fit correctly predicted 2 of the top 3 control laws that resulted in the lowest metabolic cost, and it predicted the top control law with the lowest metabolic cost 50% of the time.

#### Preference Optimization

Subject 1 completed an optimization with the objective of maximizing user comfort, instead of minimizing metabolic cost. This case study followed a similar optimization protocol, except the time spent in each control law was much shorter (23 seconds), and the participant experienced five unique control laws multiple times within a generation for a total of eleven conditions. This allowed all control laws to be directly compared with one another by asking the participant to rank the current control law as better or worse than the previous control law, and a composite “score” for each control law was used at the end of the generation to determine the parameter set of the next generation. At the end of the optimization, the parameters were as follows: stiffness = 1041 Nm/rad, peak time = 69%, rise time = 40%, fall time = 18%. In validation, in addition to comparing to the generic controller (stiffness = 900 Nm/rad, peak time = 80%, rise time = 20%, fall time = 15%), the optimization seed was also tested (stiffness = 800 Nm/rad, peak time = 65%, rise time = 30%, fall time = 20%) (Fig. 5A shows a visualization of the torque profiles). During validation, the participant was allowed to request these three controllers in any order, and was asked to rank them at the end of the trial. As shown in Fig. 5B, the participant chose the controller used as the optimization seed as the most preferred, followed by the generic controller. The optimized controller was the least preferred. We also collected the participant’s thoughts on the three controllers. He stated that the optimized controller was “too passive”, and the generic controller “too springy,” while the controller used for the optimization seed was “just right.”

**Figure 5.**
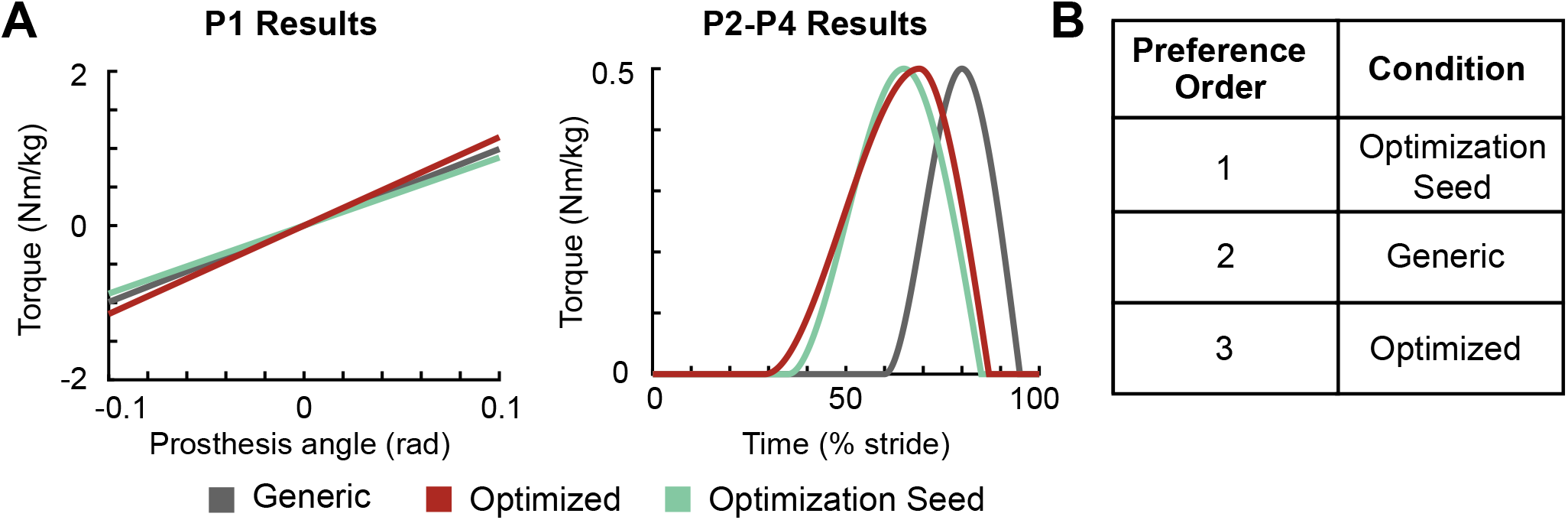
A. The parameters from all validation conditions in the 4-parameter time-based torque control optimization where preference was used as the cost function are shown. B. In validation, the controller used as the optimization seed was ranked as the most highly preferred, followed by the generic controller. The optimized controller was ranked as the least preferred.

## Discussion

We present a series of case studies in which human-in-the-loop optimization of an ankle-foot prosthesis failed to produce significant changes in metabolic cost for people with transtibial amputation. In total, five different control architectures were implemented, and between one and three participants completed the optimization protocol for any one control architecture. Mild reductions in the optimized controller of up to 3.1% compared to a generic controller were seen in some of the case studies. However, because the error in human metabolic rate measurement using indirect calorimetry has been estimated to be between 2-3%^28,29^, these reductions are not meaningful. In addition, these reductions are significantly smaller when compared to an average 24% reduction in metabolic cost as a result of optimization of unilateral ankle exoskeleton control. One case study examined an extended protocol, but the optimized controller still resulted in higher metabolic cost than during walking with a generic controller. We also examined a controller chosen by the participant via conversational tuning, but this resulted in higher metabolic rate than all comparison controllers. Finally, in one case study we attempted to optimize for preference by allowing the participant to rank controllers, but the optimized parameters were not preferred in validation.

Since we first started using the technique in 2016, human-in-the-loop optimization has been successfully used for a variety of different exoskeletons in various gait conditions and control architectures. Given its success with exoskeletons, it is surprising that HILO failed to produce meaningful changes in metabolic cost when used to tune the parameters of an ankle-foot prosthesis controller. There are several reasons why this technique may have failed, and these can be broadly separated into two categories. First, it is possible that optimization decisions that have led to successes with exoskeletons are not applicable when optimizing prosthesis control. There are many factors that can affect the optimization, such as the chosen control architecture, optimization strategy, cost function, and optimization protocol. It is possible that further modifications of these factors would lead to different results. Second, it is possible that inherent differences in user mechanics and neural control between people with amputation and those with intact limbs are the cause of the failure of HILO for prostheses. Similarly, the lack of sensory feedback, differences in learning mechanisms, or different objective functions of people with amputation could prevent HILO from being successful.

Although there are many engineering decisions that affect the results of human-in-the-loop optimization, we can hypothesize which are the most likely sources based on the case studies completed. For example, it is possible that all control architectures tested resulted in metabolic cost landscapes that were largely unvarying and did not result in minimums that were significantly lower than the cost of walking with the participants’ prescribed prostheses, even with the optimal parameters. However, we tested five different control architectures, four vastly different from each other. Both the neuromuscular and the balance controller have been successfully used in the past to reduce the metabolic cost of walking with an active prostheses below that of walking with a passive prosthesis^18,20^. In addition, the time-based torque controller is similar to one that has been successfully used to reduce the metabolic cost of walking with exoskeletons^24^. Given this information, it is unlikely that none of the control architectures tested would have been able to produce control parameters that reduce the metabolic cost of walking compared to generic control parameters, and possibly even the participant’s passive prosthesis.

Another factor that could affect the optimization is the chosen optimization protocol. The case study protocols were chosen based on the protocols successfully used for HILO with exoskeletons using a control architecture similar to the 5-parameter time-based torque controller. These previous experiments show that for a 4-parameter optimization, four generations with eight different conditions of two minutes each resulted in convergence for 9 out of 11 participants^24^. In the case studies presented here, the protocol for the 4-parameter optimization was the same, with the number of control laws for other protocols scaled based on the number of parameters. The protocol completed for the optimization of a 5-parameter controller included two additional generations. We also examined an extended protocol where the amount of time spent in each control law was increased from two to ten minutes in order to verify that the metabolic fit predicted at two minutes was similar to the average metabolic rate at the end of ten minutes. Although there were differences in metabolic rate between the two, the majority of the control laws that were predicted at two minutes to result in the lowest metabolic rate were the same as those measured at ten minutes. It is possible that an increased number of generations would produce better results, but this is unlikely because we did not observe a consistent reduction in metabolic cost or metabolic cost variation over time from one generation to the next. Nevertheless, it is possible that better results might be achieved with additional training, continued optimization or other improvements in the protocol.

The optimization strategy is an additional factor that could affect the outcome of the experiment. All pilot studies used a covariance matrix adaptation evolution strategy (CMA-ES) to calculate the next generation of control laws, which ultimately dictates the optimal control parameters chosen from the last generation. This strategy has previously been shown to be robust to measurement noise and successful in high dimensionality spaces. It is also not biased due to changes as a result of adaptation over time because it does not use information from previous generations. Other optimization strategies such as gradient descent^22,23^ or Bayesian optimization^25^ have also been used successfully for HILO of exoskeletons. However, these strategies have only been used to optimize one or two parameters simultaneously. In addition, both methods have drawbacks. Gradient descent is sensitive to measurement noise, and Bayesian optimization is not robust to human adaptation over time, as it uses all information from previous trials instead of evaluating each generation independently. It is possible, however, that using a different optimization strategy could be more effective, especially for control architectures with fewer parameters.

Finally, when defining an optimization problem it is important to consider the cost function. In this case, we must take into account two cost functions: the cost function that we choose for the optimizer and the cost function being used by the neuromuscular controller of the human in the loop as they move. The majority of prior work with HILO has simply used the minimization of metabolic rate as the cost function of the optimization, both because it is easy to quantify and because humans using exoskeletons have been shown to shown to minimize metabolic rate in real-time^30^. However, people with amputation may have additional constraints that affect their gait compared to unaffected individuals. For example, people with amputation tend to fall more frequently3and complain of socket discomfort^31^. Perhaps because of the additional importance of stability and comfort, individuals with amputation adapt gait patterns with a similar energy cost no matter what the prosthesis does. We attempted to address this issue with two case studies: one where we allowed the participant to choose their own device parameters through conversational tuning, and one where we changed the cost function of the optimization to maximize subject preference. However, the conversationally tuned controller resulted in higher metabolic cost than all other controllers tested in validation, and the controller chosen by the optimization for preference was not preferred in validation.

Independent from the factors in the optimization that can affect the results, it is possible that the contrast between the success of HILO for exoskeletons and the failure of HILO for active prostheses has to do with the differences between the groups of participants. At the joint level, people with amputation are lacking both sensing and direct control of the joint where the optimization is taking place. Perhaps one or both of these factors is necessary in order to learn during the optimization. For example, when undergoing HILO with exoskeletons, participants are able to feel the timing and magnitude of torque being applied by the exoskeletons and can adjust their gait accordingly, either to take advantage of the additional power provided by a “good” control law or to mitigate the negative affects of a “bad” control law. In contrast, people with amputation are limited in sensing to what they can feel through their socket, which has limited perception. Therefore, it is possible that a controller could be beneficial and allow them to reduce their metabolic cost if they were able to alter their gait in a certain way, but the lack of sensing prevents them from knowing how to move to best make use of the power provided by the prosthesis.

At the whole-body level, it is also possible that there are inherent differences that separate people with amputation from those with intact limbs. The additional constraints placed on people with amputation during gait has already been discussed. However, it is also possible that individuals with amputation have different learning mechanisms as well, as neural circuitry associated with motor learning could be disrupted in complex ways with the loss of a limb. Perhaps learning is possible even with the lack of sensing, but it may occur on a much longer timescale that was not captured within our experiments. Previous success in reducing metabolic cost with an active ankle-foot prosthesis has involved a long protocol in which participants are able to bring the device home with them and use it for weeks at a time before validation. One limitation of device emulators is that they are constrained to the lab, and therefore protocols of this length are not feasible. In addition, after someone has lost a limb, they must re-learn how to walk with their prosthetic limb, with prosthetists and physical therapists typically providing gait recommendations. Perhaps this training could interfere with adaptation to a new device. Or perhaps the “forced exploration” that comes from trying many diverse conditions, which is shown to be beneficial in pushing a person towards the metabolic minimum while walking with exoskeletons^32^, is actually harmful to the learning process of people with amputation.

As discussed above, there are a myriad of reasons why HILO may be more challenging when applied to powered prostheses instead of exoskeletons. By presenting the results of our case studies in which HILO did not result in meaningful changes in metabolic cost for those walking with an ankle-foot prosthesis, we do not suggest that HILO will never be successful for people with amputation. We recognize that each of our experiments contained a limited number of participants and should not be used to make broad conclusions. However, our results can be used to guide future directions for new HILO experiments with active prostheses. For example, the control architecture is a possible cause of failure, but this is less likely than other possibilities simply because we tested such a wide variety of control architectures. Our case studies also address other possible causes, such as the cost function or optimization protocol, but these are addressed less fully and warrant additional experiments.

Ultimately, HILO could still be extremely beneficial for people with amputation. Powered prostheses, if controlled properly, have the potential to reduce falls, lower metabolic cost, increase walking speed, and decrease osteoarthritis in the intact limb. Because HILO has shown that customization is important for controlling exoskeleton control parameters, it has the potential to make meaningful improvements in controlling powered prostheses as well. We hope that the results of our case studies provide a good starting point for future experiments to attempt to tease out the various causes of the failure in HILO for powered prostheses.

## Methods

### Participants

A total of five participants with unilateral transtibial amputation (N = 5, 4 male and 1 female; age = 37.8 ±14.1 [26 – 60] years; body mass = 77.8 ±8.99 [65.8 - 90.7] kg; height = 171.2 ±4.44 [167 - 178] cm; time since amputation = 13.6 ±10.4 [3 - 26] years; mean ±standard deviation) were enrolled in the case studies (Table 2). An overview of the case studies in which each participant was enrolled can be found in Table 1. All individuals provided informed consent prior to participation in the study. All study protocols were approved by the Institutional Review Board of either Carnegie Melon University or Stanford University, depending on the location at which the study took place.

### Hardware

In all case studies, subjects walked on a treadmill (Bertec, Ohio, USA) while using either an ankle-foot prosthesis emulator or their prescribed prosthesis. Walking speed for each case study was determined by participant fitness and the duration of the study (Table1). The ankle-foot prosthesis emulator consisted of off-board actuation and control hardware attached to a prosthesis end-effector (HumoTech, Pennsylvania, USA). Flexible Bowden-cable tethers transmitted mechanical power to the prosthesis. Sampling of the strain gauges and encoders of the device, as well as control commands, were implemented at 1000 Hz. All studies used one of two different ankle-foot prosthesis end-effectors (Fig. 6B). The first was a one-degree-of-freedom device with a mass of 0.96 kg capable of 41° of ankle plantarflexion, 21.7° of ankle dorsiflexion, 190 N·m of ankle plantarflexion torque, and 5 N·m of ankle dorsiflexion torque (HumoTech, Pennsylvania, USA). The second end-effector was a three-degree-of-freedom device with an actuated heel and two toe digits and a mass of 1.2 kg. This device could generate 19° of ankle plantarflexion and dorsiflexion, 140 N·m of ankle plantarflexion torque, and 100 N·m of ankle dorsiflexion torque^33^. The device used for each case study can be found in Table1.

### Metabolic Rate Calculations

To determine user metabolic rate, respirometry data was collected using a Quark CPET metabolic cart (Cosmed, California, USA). Metabolic rate was calculated using standard empirical equations^34^. In validation, net metabolic rate was calculated by subtracting standing metabolic power from the metabolic cost of all other conditions and normalizing by body mass. During optimization, most control laws were tested for two minutes of walking. As this is not typically sufficient time for metabolic rate to reach a steady-state, we fit an exponential curve to the metabolic rate and used the asymptote as the estimate of steady-state metabolic rate, as previously described^24^. One extended protocol tested each control law for ten minutes of walking.

### Optimization Trials

During optimization, participants’ metabolic rate was measured as they walked on the treadmill wearing an ankle-foot prosthesis emulator with changing control laws. A Covarience Matrix Adaptation Evolution Strategy (CMA-ES) was used to identify the control parameters, as described in Zhang et al.^24^. The goal of the optimizations was either to minimize metabolic rate or maximize subject preference. The optimization protocol, including the number of generations, number of control laws per generation, duration of each control law, walking speed, and the number of control laws tested per continuous walking bout for each case study minimizing metabolic rate is given in Table1.The protocol for the case study in which user preference was optimized is described in further detail in the Preference Optimization subsection. Prior to starting optimization trials and after every break, participants were allowed to acclimate to walking in the ankle-foot prosthesis emulator controlled using a standard spring controller. The acclimation time was one minute for the case studies using the heel stiffness controller and the 4-parameter time-based torque controller with the standard protocol, and two minutes for all other case studies.

### Validation Trials

At the end of optimization, validation trials were performed in order to compare performance of the optimized controller with one or more of the following conditions: a generic controller, the participant’s prescribed prosthesis, and a conversationally tuned controller. Following a standing baseline measurement of energy expenditure, all validation conditions were tested twice in a double reversal order to mitigate the effects of adaptation or drift. The one exception was the heel stiffness controller case study, in which the prescribed prosthesis was only tested once. The validation trials lasted 5 minutes for the case study with the heel stiffness controller, 10 minutes for the case study with the extended protocol of the 4-parameter time-based torque controller, and 6 minutes for all other case studies optimizing for metabolic rate. The metabolic rate for each trial was found by averaging the last two minutes of metabolic data. The average metabolic cost of the two validation trials per condition, normalized to user mass, determined the final metabolic cost for that condition.

### Control Parameterization

#### Heel Stiffness Controller

This case study used the three-degree-of-freedom ankle-foot prosthesis end-effector with the two toes connected to passive compression springs, and the heel being the only actively controlled digit. The stiffness of the compression springs was chosen by the participant based on comfort. Heel torque was dictated by a control architecture dependent on stiffness and damping parameters, as well as the heel angle (Fig.1A). When the heel angle is less than 0.2 rad, the heel provides a minimum torque of 1.5 N·m. However, as the heel angle increases above this threshold, stiffness and damping coefficients dictate the heel torque during loading and unloading. The resultant torque profile can be visually displayed as a parallelogram in the heel torque and angle space, with two sides of the parallelogram corresponding to the loading phase of the heel and the other two sides corresponding to the unloading phase. The bottom left point of the parallelogram is located at 0.2 rad and 1.5 N·m and is the point of transition from minimal torque behavior at a small heel angle to the higher torques at larger angles. The top right point of the parallelogram lies at a heel angle of 0.7 rad, and along a line with a slope defined by the stiffness parameter that begins at the bottom left point of the parallelogram. This stiffness parameter can vary from 50 N·m/rad to 130 N·m/rad.

While stiffness dictates the maximum torque provided at 0.7 N·m, the damping coefficient determines the trajectory the torque follows in order to reach this maximum by controlling the top left and bottom right points of the parallelogram. To find these two points, you must first determine the second axis of the parallelogram on which they occur. This second axis is determined to ensure that the heel stiffness does not exceed 130 N·m/rad at any point in the trajectory. Therefore, the first of the two points that form the second axis is found by calculating the intersection of a line with an initial value at the bottom left point of the parallelgram and a slope of 130 N·m/rad and a horizontal line with the same torque value that occurs at 0.7 radians, which varies based on the stiffness parameter. The second of these two points is found by calculating the intersection of a line with an initial value at the top right point of the parallelogram and a slope of 130 N·m/rad with a horizontal line at the minimal torque value of 1.5 N·m.

After the second axis of the parallelogram is found, the damping coefficient is used as a scaling factor that determines how far along this axis the last two points of the parallelogram are located, and is used to distinguish which points corresponds to the loading and unloading phases. A positive value of the damping coefficient indicates that the heel produces more torque in the unloading phase than the loading phase, with the bottom sides of the parallelogram corresponding to the loading phase. This results in net positive work over the gait cycle. A negative damping coefficient indicates that the heel produces less torque in the unloading phase, resulting in net negative work over the gait cycle. In addition, the magnitude of the damping coefficient is used to determine the effective “width” of the parallelogram. A damping coefficient with a value of 0 results in no width, and the torque profile becomes a function of heel angle with a given stiffness for both the the loading and unloading phases. A damping coefficient with a higher magnitude results in a greater difference between the torque magnitude in the loading phase and the torque magnitude in the unloading phase.

#### Neuromuscular Controller

The neuromuscular controller adapted the active plantarflexor component of the controller described in Eilenberg et al.^19^(Fig.2A). We varied three model parameters during optimization: *(F_max_*, **ε_ref_**, and *K_ff_*). The parameters *F_max_* and **ε_ref_** are analogous to maximum muscle isometric force and a tendon strain multiplier, respectively, with *K_ff_* acting as a feed-forward gain. Parameterizing the controller in such a way allowed for large variability in the resulting behavior of the ankle-foot prosthesis.

#### Balance Controller

We made two modifications to the balance controller previously described in Kim et al.^20^ (Fig. 3A). First, we measured deviations of center of mass velocity instead of center of mass acceleration. Center of mass velocity was determined using two string potentiometers attached to the participant’s waist and grounded to the treadmill handlebars. Since the side-to-side distance of the treadmill is known, we could calculate the center of mass of the participant with additional information from the potentiometers. By differentiating the center of mass position, we found the center of mass velocity. Second, prior work relied on an instrumented treadmill to calculate the center of mass deviation at the moment of intact limb toe-off, used to determine the ankle torque to be applied during the subsequent stance phase of the prosthetic foot. In this case study, we sampled when the prosthesis achieved foot flat due to a lack of an instrumented treadmill.

#### Time-based Torque Controller

Five parameters defined the torque profile of the time-based torque controller: device stiffness, peak torque magnitude, time to peak torque, rise time to peak torque, and fall time after peak torque. Peak time, rise time, fall time, and peak torque parameters are identical to previous work with exoskeletons24. For our work with the ankle-foot prosthesis, we also added an underlying baseline stiffness of the device (P1), creating a total of five optimization parameters (Fig. 4A). Although all parameters could be modified, three case studies utilized a controller where the peak torque was fixed at 0.5 N·m/kg, and only the other four parameters were optimized.

### Preference Optimization Protocol

Subject 1 participated in this case study focused on modifying the parameters of the 4-parameter time-based torque controller in order to maximize subject comfort, as opposed to minimizing metabolic cost. The optimization protocol included six generations with five control laws evaluated per generation. However, instead of experiencing each control law only once, the participant experienced each control law multiple times for 23 seconds each. Starting with the presentation of the second control law, the participant was asked to rate the current control law as better or worse than the previous one. Control laws were presented in an order such that all five were directly compared to each other (12345135241), and a total preference score for each control law was determined at the end of the generation. This was then used to determine the distribution of the parameter set for the next generation. Three controllers were evaluated in validation: the optimized controller, a generic controller, and a controller with the parameters used for the optimization seed. To allow for comparison of all three controllers with one another, validation consisted of 5 minutes of continuous walking, during which time the participant was allowed to request any of the three controllers at any time. At the end of the 5 minute trial, the participant ranked all three in order of preference.

## Acknowledgements

This work was funded by a National Science Foundation Graduate Research Fellowship to CGW (DGE-1656518) and National Science Foundation Grants CBET-1511177 and CMMI-1734449.

## Author contributions statement

S.H.C., V.L.C., and A.S.V. conceived the experiments, V.L.C. and A.S.V. conducted the experiments, C.G.W., A.S.V, and V.L.C analysed the results. C.G.W. wrote the manuscript. All authors edited and reviewed the manuscript.

